# Remodelled cholesteryl ester enriched lipid droplets fuel flavivirus morphogenesis

**DOI:** 10.1101/2025.06.07.658178

**Authors:** Adrianna Banducci-Karp, Sophie Brixton, Pranav N. M. Shah, Ming-Yuan Li, Georgina Fisher, Joey Riepsaame, Raman Dhaliwal, Katie L. Holden, Edward Drydale, James Bancroft, Charlotte E. Melia, Gathsaurie Neelika Malavige, Sumana Sanyal

## Abstract

Flaviviruses such as dengue and Zika viruses extensively remodel host cell membranes to create specialised replication organelles, but the role of lipid metabolism to generate them remain poorly understood.

Through systematic screens of fatty acyl transferase enzymes (MBOAT and zDHHC families) and complementary approaches including CRISPR/Cas9 gene deletions, pharmacological inhibition, proteomics, and photo-crosslinkable cholesterol analogues, we identified cholesteryl ester-enriched lipid droplets (CE-LDs) as critical host components required for flavivirus infection. CE-LD formation is mediated by Sterol O-acyltransferases 1 and 2 (SOAT1/SOAT2), whose activities were upregulated early during infection, coinciding with increased CE-LD formation and transition to liquid crystalline phases. Genetic deletion or pharmacological inhibition of either enzyme resulted in a dramatic ∼100-fold reduction in viral production. Mechanistically, CE-LDs display distinct proteomic signatures, enriched in fatty acid remodelling enzymes, GTPases, and lipid transport proteins. Photo-crosslinking experiments demonstrated direct interactions between LD-derived cholesterol and viral prM, capsid and NS1. Disrupting CE-LD formation via SOAT1/2-deficiency resulted in defective, viral RNA-free replication organelles and complete absence of immature virions. Supporting the physiological and clinical relevance of viral LD exploitation, analysis in iPSC-derived macrophages mirrored findings in Huh7 cells, and dengue patients from a Sri Lankan cohort revealed that central obesity significantly increased the risk of severe dengue haemorrhagic fever.

This study establishes CE-LDs as essential host metabolic hubs that enable flavivirus morphogenesis and identifies host LD metabolism as a promising therapeutic target for combating flavivirus infections.

---

Flaviviruses pose significant global health threats, with members such as Dengue, West Nile, and Zika affecting millions annually^1^. These mosquito-borne pathogens cause diseases ranging from mild febrile illness to severe manifestations including haemorrhagic fever, encephalitis and congenital abnormalities^1^. Despite their clinical importance, critical gaps persist in our understanding of the molecular mechanisms governing flavivirus biogenesis and host cell interactions^1^. This knowledge deficit has significantly impeded the development of effective antiviral strategies.

A hallmark of flavivirus infection is the extensive remodelling of host cell membranes to create specialised membrane compartments^1–4^. These virus-engineered compartments serve as protected factories, housing replication organelles (ROs) for viral genome replication, virion assembly sites, and transport vesicles for egress^5^, effectively shielding viral components from host immune surveillance. Biogenesis of these compartments necessitates substantial modifications in lipid metabolism, particularly involving the biosynthesis, trafficking and redistribution of lipid species to support membrane proliferation and curvature^6–10^. However, the specific host factors mediating these critical lipid modifications during flavivirus infection remain largely uncharacterised.

Central to cellular lipid homeostasis are lipid droplets (LDs), dynamic organelles consisting of a hydrophobic core of neutral lipids (primarily triacylglycerols and cholesteryl esters) surrounded by a phospholipid monolayer embedded with a complement of regulatory proteins^11,12^. Emerging evidence indicates that flaviviruses strategically manipulate both the formation and turnover of LDs during infection, suggesting these organelles serve as metabolic hubs in the viral lifecycle^6,13–16^. However, the molecular mechanisms governing flaviviruses exploitation of LDs and their transformation into specialised pro-viral structures, remain poorly understood.

We previously reported that dengue virus (DENV) infection is accompanied by early LD accumulation (10-18 hours) followed by controlled turnover via AUP1-Ube2g2 mediated lipophagy (24-48 hours), a selective autophagic pathway targeting LDs^6,13^. This temporal regulation potentially serves multiple functions: (i) supplying lipid building blocks for replication organelle biogenesis; (ii) providing fatty acids for protein acylation as a way of hijacking host factors; (iii) preventing inflammatory LD accumulation; and (iv) fueling β-oxidation to meet heightened energy demands^17^.

In this study, we demonstrate that flavivirus infection transforms LDs into specialised cholesteryl ester-enriched structures (CE-LDs) with liquid crystalline phases and extensively remodelled proteomes. LD-alterations were accompanied by an increase in both lipid and protein fatty acylation in Zika virus (ZIKV) infected cells. Through systematic screens of fatty acyl transferase enzymes (MBOAT^18^ and zDHHC families^19^) we identified Sterol O-acyltransferase 1 and 2 (SOAT1 and SOAT2) as critical host enzymes mediating this transformation. Also known as Acyl-CoA:cholesterol acyltransferases (ACAT1 and ACAT2), these enzymes serve as key regulators of cellular cholesterol metabolism^20^. These are ER-resident integral membrane proteins that catalyse the formation of cholesteryl esters from long-chain fatty acyl-CoA and cholesterol^21^. This esterification process generates transport-competent forms of cholesterol that are important for cholesterol homeostasis and LD formation^22^.

Our findings reveal that SOAT1/2-mediated CE-LD formation creates liquid crystalline metabolic hubs enriched in fatty acid remodelling enzymes, Rab-GTPases, and lipid transport proteins that are essential for flavivirus morphogenesis. Photo-crosslinking experiments demonstrated direct interactions between LD-derived cholesterol and viral structural proteins (prM, capsid) and the non-structural protein NS1. Disruption of SOAT1/2 activity resulted in defective, viral RNA-free replication organelles and complete absence of immature virions, both in Huh7 cells and iPSC-derived macrophages. These mechanistic insights are clinically relevant, as analysis of dengue patients from a Sri Lankan cohort revealed that central obesity significantly increases severe disease risk, likely reflecting enhanced viral exploitation of CE-LDs. This study establishes CE-LDs as essential host components for flavivirus infection and identifies LDs as a source of potential therapeutic targets.

## Results

### Lipid droplet dynamics in live and fixed cells during ZIKV infection

We previously characterised Aup1/Ube2g2-dependent lipid droplet (LD) turnover during dengue infection^6,13^. Building on these observations, we investigated the temporal dynamics of LDs during ZIKV infection using live-cell imaging. We infected Huh7 cells with ZIKV at different MOIs and monitored LD dynamics in real-time through fluorescence microscopy (**Figure 1A**). Live cell tracking revealed distinct temporal patterns of LD accumulation in ZIKV-infected cells, with migration of LDs to perinuclear, dsRNA-positive regions, followed by their turnover (**Figure 1A**). Quantitative analysis of LD numbers demonstrated infection-dependent alterations in LD dynamics, with an initial increase followed by a decrease in LD abundance over the 24-hour period (**Figure 1B**).

**Figure 1.**
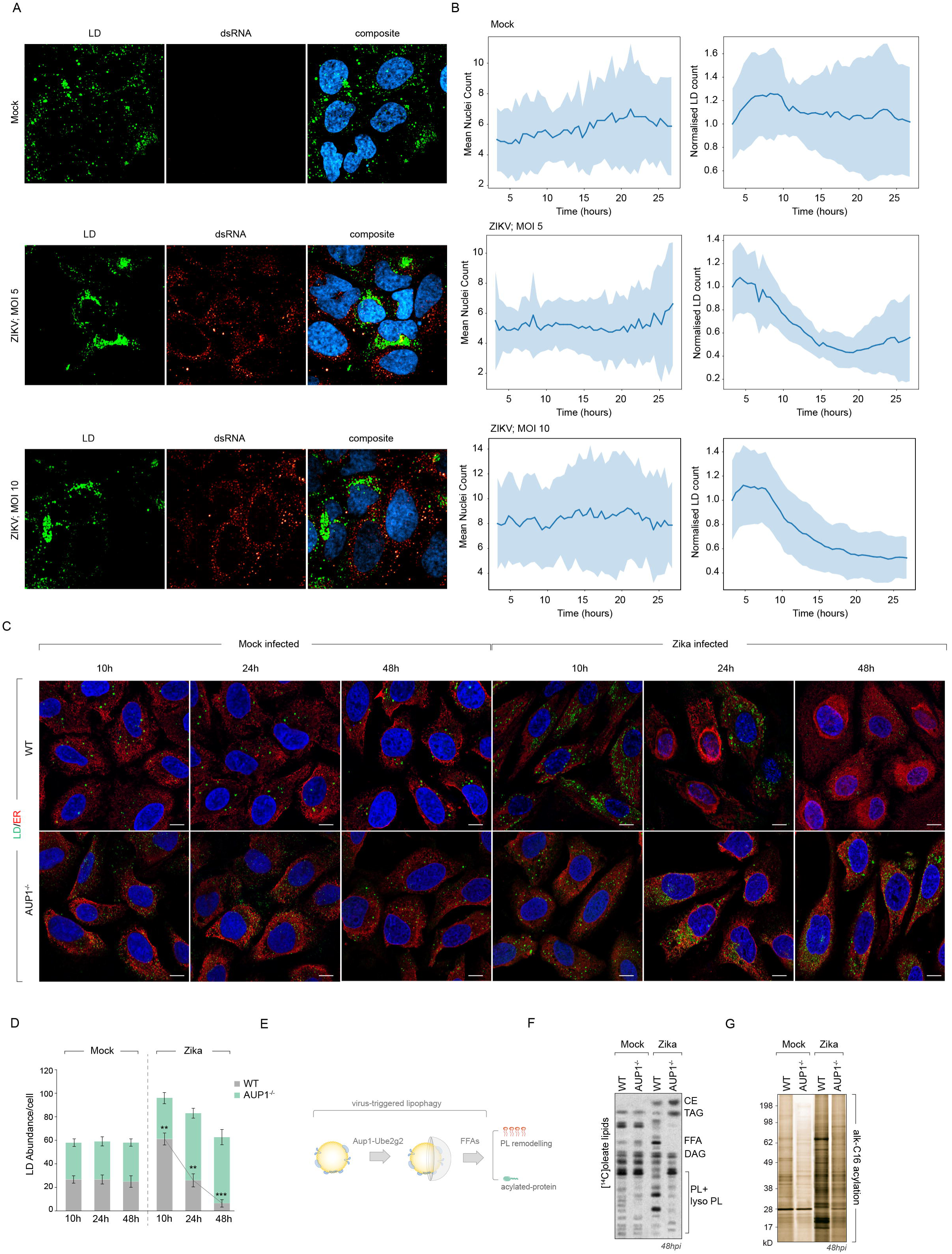
Flavivirus infection alters lipid droplet dynamics. **(A)** Maximum projections at 24h from live-cell microscopy of ZIKV-infected Huh7. LDs (green, BODIPY 493/503) and viral dsRNA (red). **(B)** Quantitative temporal analysis of LD abundance per cell derived from live-imaging experiments at different multiplicities of infection (MOI 5 and MOI 10). Data represent mean ± SEM from n≥50 cells per condition with statistical analysis performed using time-series regression modeling. **(C)** Confocal microscopy of WT and AUP1^-/-^ Huh7 cells stained with BODIPY (LD, green), Sec61 (ER, red) and DAPI (nuclei, blue) at 10, 24, and 48h in control and ZIKV-infected cells. **(D)** Quantification of LD numbers per cell in WT and Aup1^-/-^ cells over infection time course. n≥30 cells per condition across 3 independent experiments; data show mean ± SEM. **(E)** Schematic of AUP1-Ube2g2 mediated lipophagy releasing fatty acids for enhanced acylation and phospholipid remodelling during infection. **(F-G)** Acylation profiling by TLC and silver staining in mock vs. ZIKV-infected WT and Aup1^-/-^ cells at 48h p.i. **(F)** Lipid acylation in [^14^C]oleate labelled cells and **(G)** protein acylation levels in alkyne C16 labelled cells subjected to biotinylation by click reaction.

To complement these live-imaging studies, we performed fixed-cell immunofluorescence analysis at discrete time points (10h, 24h, and 48h post-infection) in both mock-infected and ZIKV-infected cells (**Figure 1C**). This approach confirmed the characteristic temporal pattern of early LD accumulation followed by gradual turnover, consistent with our previous observations in DENV infection^13^. In contrast, Aup1^-/-^ cells displayed impaired LD turnover, maintaining elevated LD numbers throughout the infection time course (**Figure 1C**, **1D**).

Given that LD turnover releases free fatty acids, we hypothesised that this process might enhance lipid and/or protein acylation during ZIKV infection (**Figure 1E**). Comprehensive acylation profiling revealed significant increases in both lipid and protein acylation in WT cells (**Figure 1F**, **1G**), which was inhibited in Aup1^-/-^ cells. These findings establish a link between LD turnover and enhanced acylation during infection (**Figure 1F**, **1G**).

To identify the specific acyltransferases mediating these modifications, we conducted parallel targeted screens of two enzyme families in ZIKV-infected cells: the ZDHHC S- palmitoyltransferases (**Figure 2A**) and the membrane-bound O-acyltransferases (MBOATs) (**Figure 2B**) in ZIKV infected cells. The zDHHC family members primarily mediate protein S-palmitoylation and have established roles in SARS-CoV-2 infection and immunomodulation^23–26^. MBOATs catalyse fatty acid transfer to diverse substrates including lipids and proteins, with several members previously implicated in viral replication^27^.

**Figure 2.**
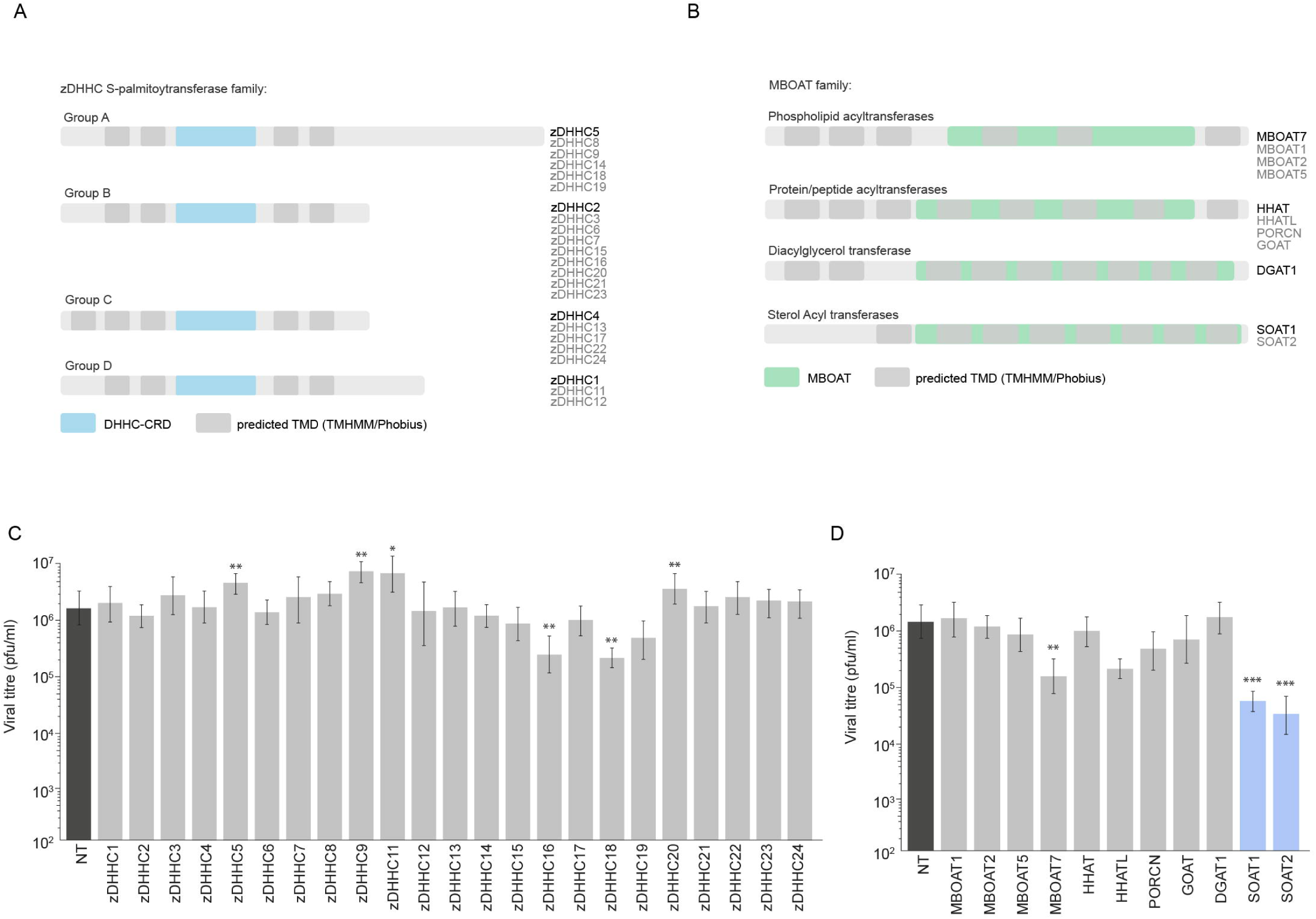
Flavivirus infection alters lipid droplet dynamics. **(A-B)** Schematic of ZDHHC palmitoyltransferases and MBOAT acyltransferases (11 family members). **(C-D)** Viral titres (pfu/ml) at 24h p.i. following siRNA knockdown of ZDHHC family and MBOAT family members. n=3 replicates; **p<0.01, ***p<0.001 vs. scrambled control.

To determine the functional significance of these acyltransferases during ZIKV infection, we performed siRNA-mediated gene silencing against individual MBOAT and ZDHHC family members. Knockdown efficiency was confirmed by mRNA levels (**Figure S1A and S1B**). While depleting several zDHHC proteins increased virus production (**Figure 2C**), depleting SOAT1 and SOAT2 (enzymes involved in cholesteryl ester synthesis) substantially reduced virus production (**Figure 2D**). In contrast, depleting DGAT1 (involved in triacylglycerol synthesis) had no detectable impact on viral titres (**Figure 2D**). These results indicate that SOAT1 and SOAT2, which specialise in cholesterol esterification, play important roles in supporting efficient ZIKV replication.

### SOAT1 and SOAT2 activities are required for productive viral infection

SOAT1 and SOAT2 (alternatively known as ACAT1 and ACAT2) are ER-resident cholesterol acyltransferases that convert free cholesterol to esterified cholesterol to be deposited into LDs^28^ (**Figure 3A**). To investigate their potential involvement in flavivirus infection, we first examined their RNA expression profiles and activity during virus infection (**Figure 3B**). ZIKV infection significantly upregulated SOAT2 mRNA levels, while SOAT1 mRNA levels showed less pronounced increase that occurred only at 24 hours post infection. In contrast, DGAT1 and DGAT2 mRNA levels remained relatively stable, with DGAT2 mRNA levels significantly decreasing at 48 hours post infection. Consistent with increased SOAT1 and SOAT2 activity, virus-infected cells contained more cholesterol esters (**Figure 3C**).

**Figure 3.**
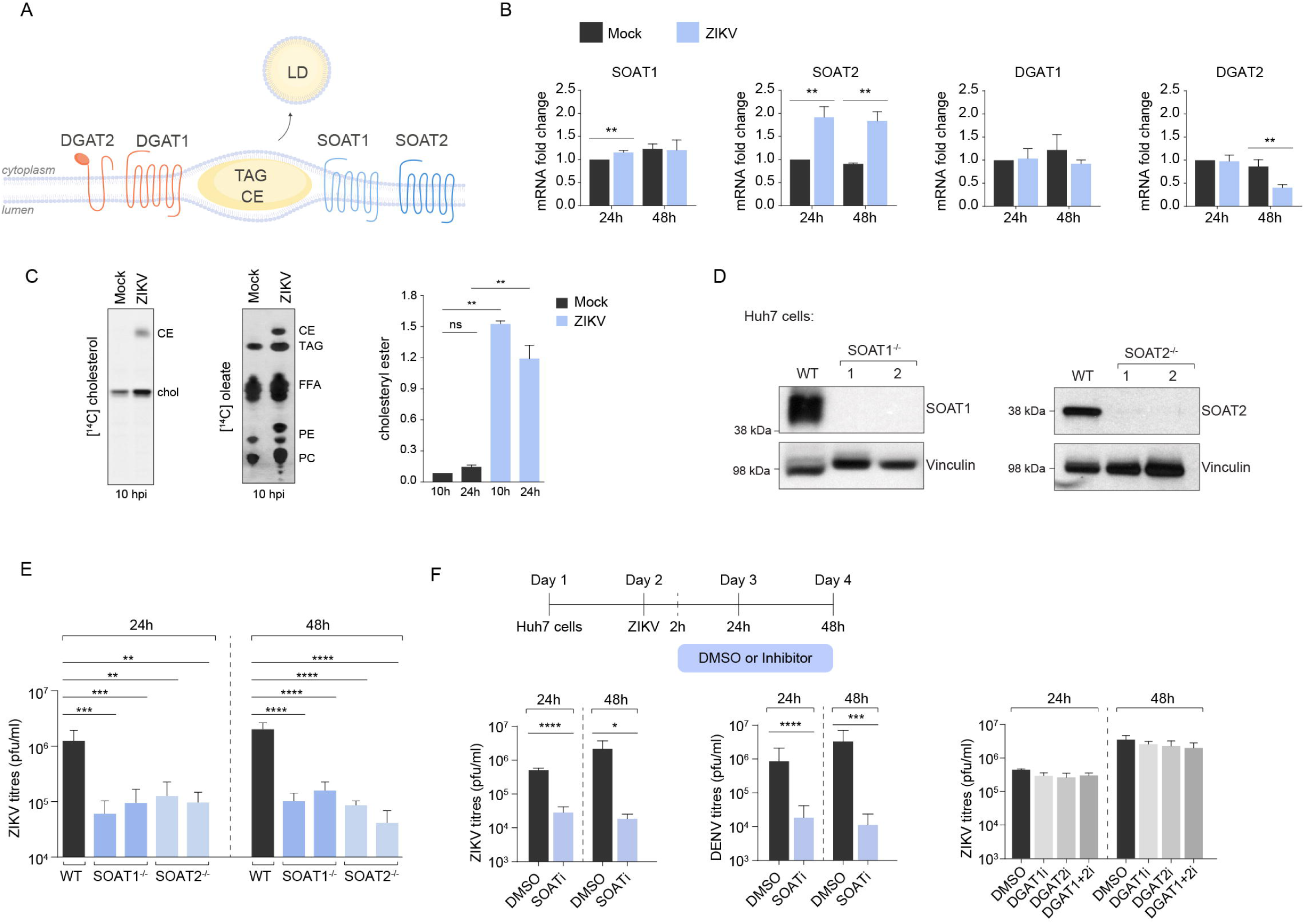
SOAT1 and SOAT2 activities are required for productive viral infection. **(A)** Schematic of LD biogenesis showing SOAT1/2-mediated conversion of free cholesterol and fatty acyl-CoA to cholesteryl esters (CE) and DGAT1/2-mediated synthesis of triacylglycerols (TAG) for LD formation. **(B)** qRT-PCR analysis of acyltransferase mRNA expression in Huh7 cells at 24h and 48h post-ZIKV infection (MOI 2). Data normalised to beta-actin; n=3 independent biological replicates; *p<0.05 vs. mock. **(C)** Cholesteryl esterification activity assay using [^14^C]-cholesterol (left) and [^14^C]-oleate incorporation in mock and ZIKV-infected cells. Lipids are extracted from cells at 10 hpi and resolved by TLC. Percentage of cholesteryl esters is measured by densitometry at 10h and 24h post infection. **(D)** Western blot validation of CRISPR/Cas9-generated SOAT1^-/-^ and SOAT2^-/-^ Huh7 cell lines. Vinculin is used as loading control. **(E)** Viral titres (PFU/ml) in culture supernatants from WT, SOAT1^-/-^, and SOAT2^-/-^ cells at 24h and 48h post-ZIKV infection. n=4 independent experiments; ***p<0.001 vs. WT. **(F)** Viral titres following pharmacological inhibition. Left panels: Avasimibe (20µM) treatment at 2h post infection in Huh7 cells infected with ZIKV compared to DMSO (n=3). Middle panel: Avasimibe (20µM) treatment in Huh7 cells after DENV infection, at 24h and 48h (n=3). Right panel: Individual and combined inhibition of DGAT inhibitor vs DMSO controls in ZIKV infected cells.

To assess the functional impact of SOAT1 and SOAT2 on Zika virus production, we generated Huh7 cells with deletions in these genes via CRISPR/Cas9 gene-editing (**Figure 3D**). Confocal microscopy revealed a marked decrease in LD numbers in the SOAT-deficient cells compared to wild-type cells (**Figure S2A**), confirming the expected disruption of cholesteryl ester synthesis and storage. SOAT1^-/-^ and SOAT2^-/-^ cells dramatically reduced virus production by ∼100-fold at 24 and 48 hours post-infection (**Figure 3E**). To validate these findings through an independent approach, we treated wild-type cells with Avasimibe, a specific inhibitor of SOAT1/2 enzymatic activity^29^ (**Figure 3F**). Increasing concentrations of Avasimibe similarly reduced viral titres in supernatants at 24h and 48h post-infection for both ZIKV and DENV without causing cytotoxicity at these concentrations (**Figure 3F, Figure S2B**). This is in line with a previous report showing reduced virus production in a range of cells and tissues upon treatment with a SOAT1 inhibitor^30^. In contrast, inhibiting DGAT1/2 had no measurable impact on virus production (**Figure 3F**).

### SOAT1 and SOAT2 activities are required for morphogenesis of virions

To determine the mechanism underlying impaired virus production in SOAT1^-/-^ and SOAT2^-/-^ cells, we visualised dsRNA (marking virus replication organelles) and viral E-protein (indicating viral assembly sites) in infected wild-type, SOAT1^-/-^ and SOAT2^-/-^ Huh7 cells (**Figure 4A**). Immunofluorescence microscopy revealed that SOAT1 or SOAT2 deletion substantially reduced dsRNA abundance and caused an even more profound decrease in E-protein clusters (**Figure 4A**). We confirmed this reduction through RT qPCR quantitation of viral RNA (**Figure 4B**) and immunoblotting for viral proteins (**Figure S3A**). Interestingly, SOAT1/2 inhibitor treatment substantially decreased structural proteins and NS1 levels (**Figure S3A, S3B**), while viral RNA levels remained unchanged, suggesting that these enzymes do not directly affect NS5-dependent replication (**Figure S3C**). To test whether virus particle assembly is also affected, we employed our previously established virus-like-particle system expressing prME alone (**Figure 4C**) to measure structural protein levels^14^ independent of replication. Similar to live infection, SOAT1^-/-^ and SOAT2^-/-^ cells had profoundly reduced prM and E levels (**Figure 4D**).

**Figure 4.**
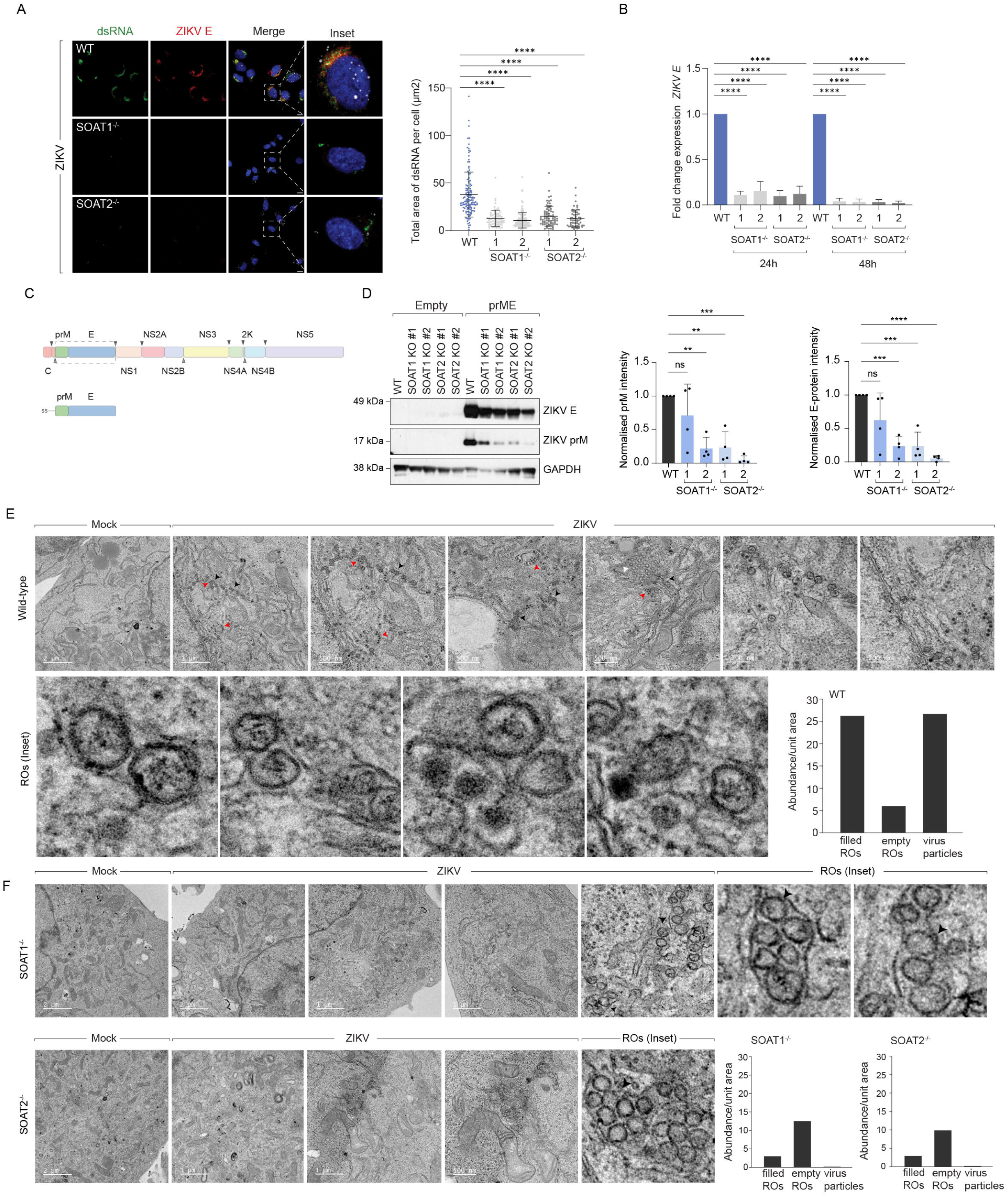
SOAT1 and SOAT2 activities are required for morphogenesis of virions. **(A)** Immunofluorescence microscopy of WT, SOAT1^-/-^, and SOAT2^-/-^ Huh7 cells at 24h post- ZIKV infection. dsRNA (green) marks viral replication organelles, ZIKV E protein (red) indicates assembly sites, DAPI (blue) shows nuclei. LDs are stained with BODIPY 493/503 in white. Insets show higher magnification of boxed regions. Scale bars 10µm. Right panel: Quantitative analysis from panel A. Total area of dsRNA per cell (μm²). n≥50 cells per condition across 4 independent experiments; ****p<0.0001. **(B)** Fold change in ZIKV E RNA expression normalised to WT controls. N=3 independent experiments; ****p<0.0001. **(C)** Schematic of virus-like particle (VLP) system expressing flavivirus structural proteins prM and E for assembly analysis independent of replication. **(D)** Western blot analysis and quantification of prM and E protein levels in VLP-transfected cells. Left: Representative blot showing prM (∼17kDa), ZIKV E (∼50kDa), and GAPDH loading control. Right: Normalised prM and E protein intensity in WT vs. SOAT knockout cells. n=3 biological replicates; **p<0.01, ***p<0.001. **(E)** Transmission electron microscopy of WT Huh7 cells at 24h post-ZIKV infection (MOI 2). Top row: Mock and infected cells. Bottom row: Magnified ZIKV- infected cells showing expanded ER with vesicle packets (replication organelles, red arrows) containing electron-dense viral RNA adjacent to assembled virions. Right: Quantification of indicated structures per field. n=20 fields per condition. **(F)** TEM analysis of SOAT1^-/-^ and SOAT2^-/-^ cells showing dramatically reduced replication organelle formation and complete absence of virus particles. Replication organelles present are predominantly empty (devoid of electron-dense material). Right: Quantification showing reduced RO abundance and absent virus particles in knockout cells compared to WT.

To determine whether SOAT1/2-deficiency impacted viral RNA stability and virion assembly on account of defective replication organelle and assembly sites, we used transmission electron microscopy on chemically fixed and resin-embedded sections to assess ultrastructural defects (**Figure 4E**). Wild-type virus-infected cells frequently displayed expanded ER containing vesicle packets and immature virus particles near the vesicle packets. These replication organelles contained electron-dense material, likely viral RNA, positioned adjacent to assembled virions in the wild-type infected cells (**Figure 4E**). In contrast, SOAT1^-/-^ or SOAT2^-/-^ cells had substantially reduced numbers of replication organelles, and those present predominantly lacked electron-dense material (highlighted in inset) (**Figure 4F**). Most significantly, we could not detect any assembled virions in SOAT1^-/-^ or SOAT2^-/-^ cells (**Figure 4F**). Similarly, SOAT-inhibitor treatment impaired biogenesis of replication organelles and viral progeny (**Figure S3D**). Inhibitor-treated cells also displayed defective mitochondrial morphology, indicating potentially altered β-oxidation. Collectively, these data indicate that SOAT1 or SOAT2 deficiency results in defective replication organelles and assembly sites, likely causing degradation of viral RNA and abolishing assembly of viral progenies.

### Remodelled cholesteryl ester enriched LDs supply lipids for ROs and virus assembly sites

SOAT1 and SOAT2 convert long-chain fatty acids and free cholesterol into cholesteryl esters that are deposited into LDs. ZIKV infection induces SOAT1 and SOAT2, increasing the amount of cholesteryl esters (**Figure 3B**, **3C**). Consistent with this, TEM analyses revealed striking morphological differences between LDs in uninfected versus ZIKV-infected cells indicative of a change in composition (**Figure 5A**). Interestingly, LDs in virus-infected cells were mostly ER-enwrapped. When visualised in FIB-milled samples, LDs in virus-infected cells frequently displayed a liquid crystalline phase (with 3.90 nm spacing of the lattice), a characteristic feature of cholesteryl ester-enriched LDs^31,32^ (**Figure 5B**, **5C**).

**Figure 5.**
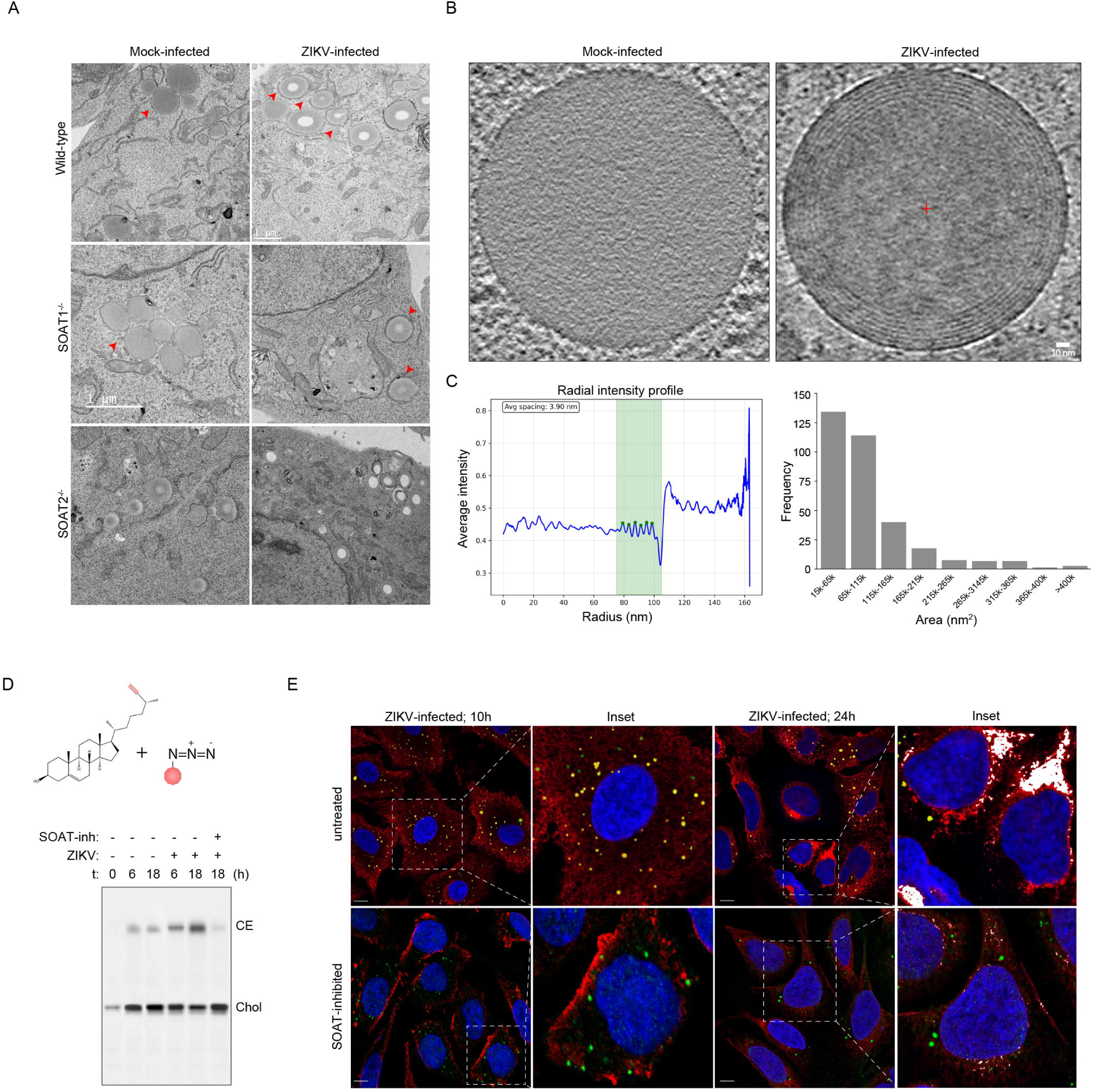
Remodelled cholesteryl ester enriched lipid droplets supply lipids for replication organelles and virus assembly sites. **(A)** Transmission electron microscopy of lipid droplets (red arrows) in mock-infected and ZIKV-infected Huh7 cells (24h p.i.) across WT, SOAT1^-/-^, and SOAT2^-/-^ cell lines, showing ER-enwrapped LDs with altered LD morphology in infected WT cells. **(B)** cryo-EM of individual lipid droplets from FIB-milled samples showing internal structure. Mock-infected cells display homogeneous LD structure while ZIKV-infected cells exhibit liquid crystalline phases characteristic of cholesteryl ester-enriched LDs. **(C)** Quantitative analysis of LD internal organisation. Left: Radial intensity profile analysis measuring structural organisation within LDs (ring spacing ∼3.9nm indicates liquid crystalline phase). Right: Histogram showing distribution of LD areas (nm²). **(D)** Chemical biology approach using cholesterol alkyne analogue for metabolic labelling. *Top*: Structure of alkyne cholesterol probe. *Bottom*: Time course of labelling followed by ZIKV infection -/+ SOAT inhibition. TLC analysis for resolving free cholesterol (Chol) and cholesteryl esters (CE). **(E)** Temporal tracking of cholesterol localisation during ZIKV infection using click chemistry-labelled alkyne cholesterol. Confocal microscopy at 10h and 24h p.i. in untreated and SOAT inhibitor-treated cells. DAPI (blue) shows nuclei, cholesterol signal (red) demonstrates redistribution from LDs (green) to viral replication/assembly sites (dsRNA, white) over time. Insets show higher magnification of boxed regions. Scale bars 10µm.

To directly investigate this compositional remodelling of LDs, we applied a chemical biology approach using a cholesterol alkyne analogue^33^. This probe is metabolically incorporated into cells, esterified and deposited into LDs. Through click chemistry, we attached a fluorescent tag onto alkyne cholesterol to visualise its subcellular localisation in virus-infected wild-type versus SOAT-deficient cells (**Figure 5D**). TLC analysis confirmed that infected cells displayed significantly higher conversion of free cholesterol to cholesteryl esters compared to mock-infected cells, in a SOAT-dependent manner (**Figure 5D**). Temporal tracking of cholesterol localisation revealed a dynamic redistribution pattern during infection (**Figure 5E**). At early timepoints, cholesteryl esters predominantly accumulated within LDs. However, as infection progressed, this cholesterol redistributed to viral replication and assembly sites, where it colocalised with dsRNA (**Figure 5E**). In contrast, SOAT1/2 inhibition caused cholesterol to accumulate in the plasma membrane and prevented co-localisation with replication organelles during infection (**Figure 5E**).

To identify the viral proteins that directly interact with cholesterol, we employed a bifunctional cholesterol analogue featuring both photo-crosslinking capability (via a diazirine group) and click chemistry reactivity (via an alkyne group)^34^. We metabolically labelled mock and ZIKV-infected cells with this cholesterol analogue and exposed them to UV-light to activate the photo-crosslinking group, followed by Cu-catalysed click reaction with azide-biotin (**Figure 6A, B**). Streptavidin pulldown and silver staining confirmed capture of cross- linked proteins in both mock and virus-infected cells, but with distinct profiles (**Figure 6C**). Immunoblotting revealed that viral proteins prM, capsid and NS1 interacted with cholesterol (**Figure 6D**), in line with previous reports^35,36^. Importantly, SOAT1/2 inhibitor treatment abolished these interactions, indicating that LD-derived cholesteryl esters – rather than free cholesterol – represent the primary source of cholesterol for viral protein interactions (**Figure 6D**). To experimentally confirm this, we treated mock and infected cells with lipolysis inhibitors that target ATGL^37^, HSL/MAGL^37^ and NCEH1^38^ (**Figure 6E**). Both orlistat, (broad lipase inhibitor that also blocks FASN)^39^, and JW480 (a selective neutral cholesteryl ester hydrolase inhibitor)^38^ substantially reduced virus production (≥1-log) at 24h post infection (**Figure 6F**), supporting our hypothesis. The modest effect of ATGL inhibition (NG-497) likely reflects partial block in lipophagy of large LDs^40^.

**Figure 6.**
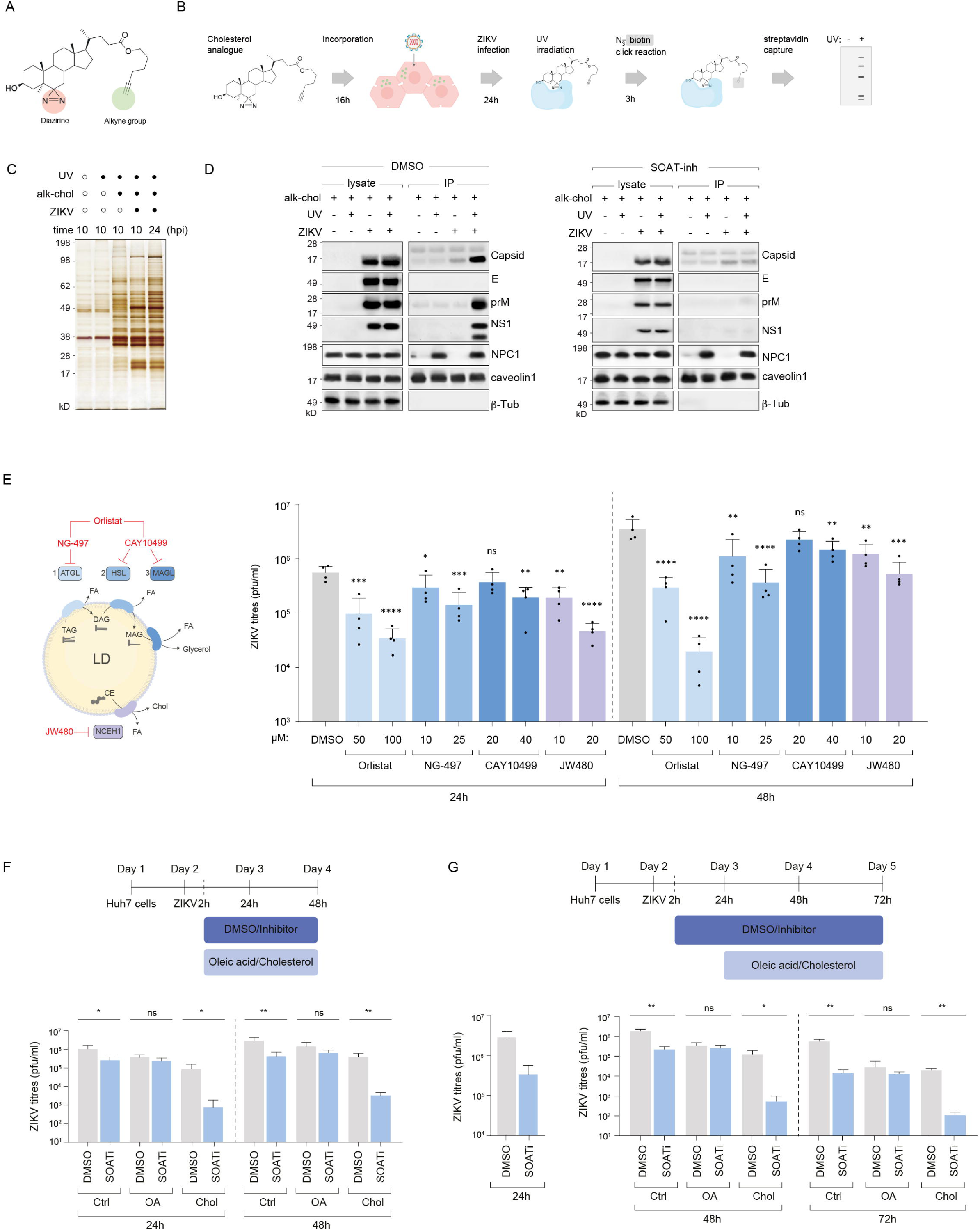
Cholesteryl ester-derived cholesterol directly interacts with viral proteins and cannot be substituted by free cholesterol. **(A)** Structure of bifunctional cholesterol analogue containing both photo-crosslinking capability (diazirine group) and click chemistry reactivity (alkyne group) for identifying cholesterol-protein interactions. **(B)** Experimental workflow for photo-crosslinking analysis. Cells are metabolically labelled with cholesterol analogue (16h), infected with ZIKV (24h), UV-irradiated for photo-crosslinking (3h), followed by Cu-catalysed azide-biotin click reaction and streptavidin capture. **(C)** Silver staining of streptavidin-captured proteins showing distinct profiles between mock and ZIKV-infected cells with different experimental conditions in a UV exposure and alkyne-cholesterol (alk-chol) dependent manner. Distinct protein profiles are captured between mock and ZIKV infection,10h and 24h post-infection. **(D)** Western blot analysis of cholesterol-crosslinked proteins from streptavidin captured fractions in control DMSO-treated cells (left) and SOAT inhibitor treated cells (right). Caveolin and NPC1 serve as positive controls and β-tubulin serves as the loading control. **(E)** Schematic of LD hydrolysis inhibition with NG-497 (ATGL), CAY10499 (non-selective lipase; inhibits HSL and MAGL), orlistat (broad lipase inhibitor that also covalently blocks the FASN thioesterase), and JW480 (NCEH1; neutral cholesteryl-ester hydrolase). **(F)** ZIKV titres (pfu/ml) measured at 24h and 48h post-infection for indicated compounds and concentrations (µM); DMSO serves as vehicle control. Error bars represent mean±SD; **p<0.01, ***p<0.001, ****p<0.0001 by one-way ANOVA with Dunnett’s correction. **(G-H)** Time-of-addition rescue experiments with oleic acid or free cholesterol in SOAT-inhibited cells. Experimental timeline showing SOAT inhibitor treatment with oleic acid (OA) or cholesterol (Chol) supplementation at the start of or 24h post infection. Viral titres (PFU/ml) measured in supernatants from control and lipid supplemented cells by plaque assays. (H) *left panel*: supernatant collected before lipid supplementation at 24h to confirm reduction in viral titres upon SOAT inhibition. n>3 biological replicates; *p<0.05, **p<0.01, ns=not significant vs. DMSO control.

To test whether exogenous lipids could rescue viral titres in SOAT1/2-inhibited cells, we supplemented either free cholesterol or fatty acids in the form of oleic acid in time-of-addition experiments (**Figure 6G**, **6H**). When we added oleic acid to SOAT1/2-inhibited cells, virus production was partially rescued compared to DMSO-treated cells. In contrast, addition of free cholesterol – either at the same time of inhibitor addition, or 24 hours after inhibition addition – exacerbated the difference in viral titres between DMSO and SOAT-inhibited conditions regardless of timing (**Figure 6G, 6H**). This indicates that the specific LD remodelling and lipid mobilisation during infection are crucial for ER-reorganisation required for productive virus morphogenesis, while free cholesterol can be detrimental to virus production.

### Virus-induced LD remodelling creates metabolically active organelles essential for morphogenesis and disease pathogenesis

To determine the molecular mechanisms by which cholesteryl ester rich (CE)-LDs support membrane remodelling during flavivirus infection, we isolated LDs on discontinuous sucrose gradients using established protocols^41,42^. LD fractions showed appropriate marker enrichment, consistent with high purity (**Figure 7A**). Compared to mock controls, ZIKV-infection dramatically altered the LD proteome, enriching fatty acid remodelling enzymes, Rab-GTPases, multiple lipid transport proteins (LTPs) including the class of bridge-like LTPs, and sphingomyelinases (**Figure 7B**). In addition, infection enriched ER scramblases TMEM41B/VMP1 (regulators of autophagy, LD biogenesis and phospholipid equilibration), indicating that they likely interact with these CE-rich LDs. These alterations collectively suggest a virus-induced remodelling of LDs at both the lipidome and proteome levels, transforming them into metabolically active hubs optimised for lipid mobilisation and delivery. We examined key host proteins in lysates from infected cells. Remarkably, genetic deletion of either SOAT1 or SOAT2 completely abolished expression of specific proteins (data not shown) such as TMEM41B (a lipid scramblase previously implicated in flavivirus replication)^43,44^. This loss was recapitulated in SOAT1/2-inhibited cells (**Figure 7C**). These findings suggest a pathway in which SOAT1/2 enzymatic activity generates specialised CE- LDs with modified proteomes that include critical lipid transport machinery, such as LTPs and the scramblase TMEM41B. This remodelling most likely occurs at ER-LD contact sites, where regulated hydrolysis and lipid remodelling generate the lipid species necessary for virus morphogenesis. To test whether SOAT activity drives organelle partitioning, we fractionated organelles ± SOAT inhibitor and probed for key viral and host proteins^5,14^. All of the factors were reduced or redistributed when SOAT was inhibited (**Figure 7D**). Collectively, our study demonstrates that flaviviruses, particularly DENV and ZIKV induce extensive reprogramming of LDs that promotes virus production. These findings have important implications for understanding severe disease manifestations such as liver damage in infected individuals with obesity and related metabolic disorders^45^.

**Figure 7.**
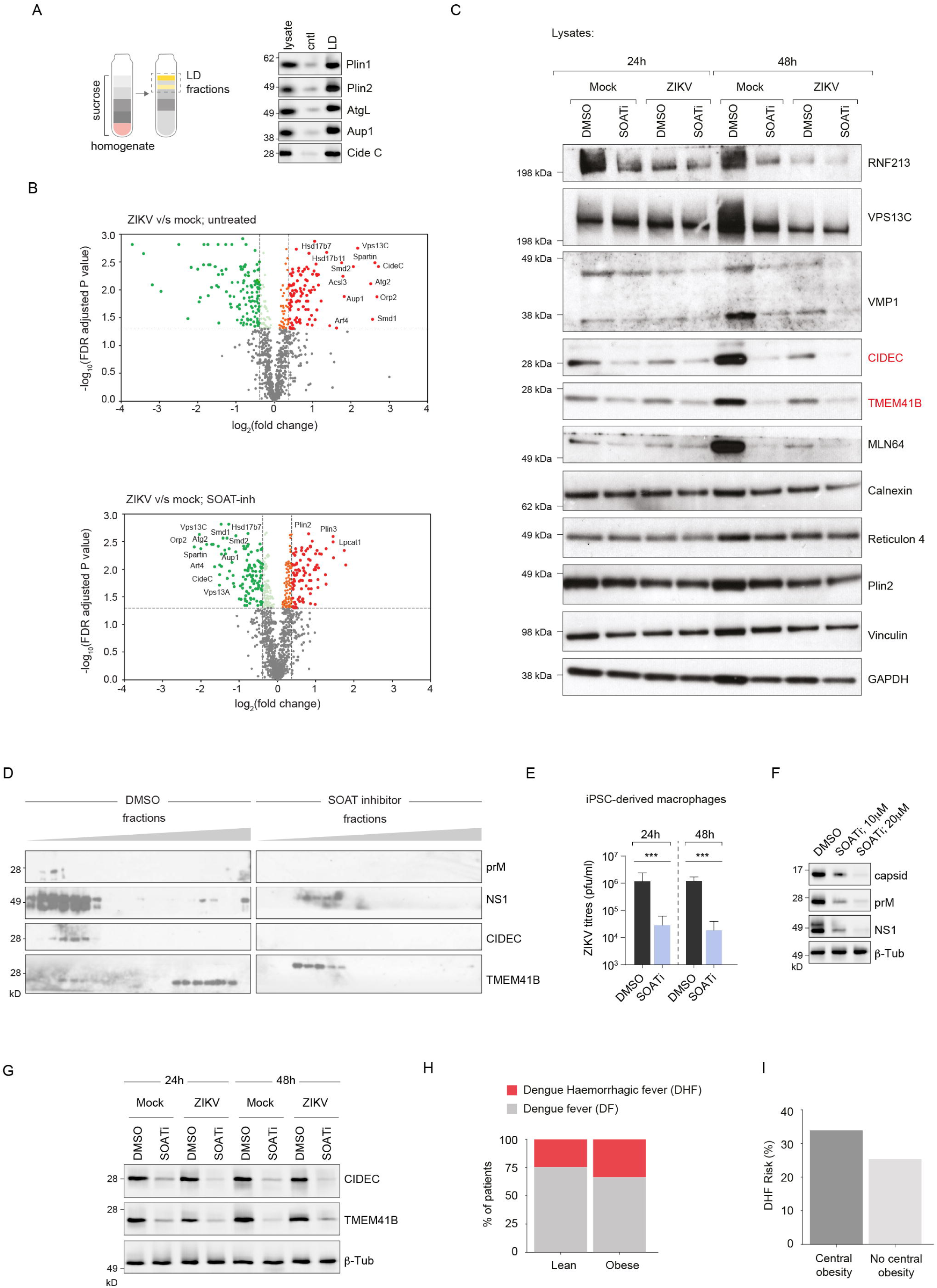
Cholesteryl ester-rich LDs are enriched in fatty acid remodelling and lipid transport proteins. **(A)** Schematic of lipid droplet isolation protocol using sucrose density gradient centrifugation for proteomic analysis of LD fractions from mock and ZIKV-infected cells; *(right panel)* enrichment of LD markers. **(B)** Volcano plots of LD proteome analysis comparing ZIKV-infected vs. mock cells. Top: Untreated conditions with significant enrichment of fatty acid remodelling enzymes, Rab-GTPases, lipid transport proteins (LTPs), and sphingomyelinases (red dots = upregulated, green dots = downregulated). Bottom: SOAT inhibitor-treated conditions with altered protein recruitment patterns. FDR-adjusted p-values plotted against logL(fold change); dotted line indicates significance threshold. **(C)** Western blot validation of key ER and membrane remodelling proteins. Analysis of whole cell lysates at 24h and 48h post-infection comparing mock vs. ZIKV infection under DMSO control or SOAT inhibitor treatment. Proteins indicated: CIDEC (lipid transferase), TMEM41B (lipid scramblase), VMP1, and MLN64, Calnexin, Reticulon 4 and Plin2. Vinculin, and GAPDH serve as loading controls. **(D**) Organelle-fractionation followed by immunoblotting in infected cells ± SOAT inhibitor. Indicated proteins are viral prM and NS1 and host CIDEC and TMEM41B; SOAT inhibition reduces LD association/redistribution of these proteins. **(E)** Human iPSC-derived macrophages: extracellular ZIKV titres (pfu/ml) at 24 h and 48 h ± SOAT inhibitor. Bars show mean ± s.e.m; statistics by one-way ANOVA with Dunnett’s test vs DMSO. **(F-G)** iPSC- macrophage lysate blots for viral proteins (capsid, prM, NS1) (F) and host proteins Cide C and TMEM41B (G) under DMSO or SOAT-inhibitor (10-20 μM). **(H)** Disease severity distribution by central obesity status. Stacked bar chart showing the proportion of dengue fever (DF, gray) and dengue haemorrhagic fever (DHF, red) in lean (n=303) and centrally obese (n=258) patients. Central obesity was defined as waist circumference ≥80 cm in women or ≥90 cm in men. p=0.0407, Fisher’s exact test. **(I)** DHF risk comparison by central obesity status. Bar chart displaying the percentage risk of developing dengue haemorrhagic fever in patients with central obesity (33.3%) versus those without central obesity (25.4%). The absolute risk difference of 7.9 percentage points represents a significant increase in severe disease risk associated with central obesity (relative risk 1.22, 95% CI: 1.01-1.46; odds ratio 1.47, 95% CI: 1.02-2.12).

To connect these mechanisms to a clinically relevant human lineage, we tested iPSC-derived macrophages, a recognised model for ZIKV/DENV infection^46^. SOAT inhibition reduced infectious titres at 24–48 h (**Figure 7E**) and diminished prM/NS1 abundance alongside CIDEC/TMEM41B in immunoblots (**Figure 7F-G**), mirroring our findings in Huh7 cells. These results support a cell-type independent requirement for CE-LD mobilisation and remodelling in morphogenesis.

Given that central obesity elevates LD burden and alters cholesterol handling, we examined a Sri Lankan dengue cohort (561 patients) stratified by central obesity status, defined by waist-circumference cut-offs widely used for Asian populations (≥90 cm men; ≥80 cm women)^47^. Analysis of disease severity distribution revealed a clear association between central obesity and severe dengue manifestations (**Figure 7H**). While both lean and obese patient groups showed predominantly dengue fever (DF) cases, obese patients demonstrated a markedly higher proportion of dengue haemorrhagic fever (DHF) compared to lean patients. This difference in disease severity distribution was statistically significant (p=0.0407, Fisher’s exact test). Quantitative analysis of DHF risk showed that patients with central obesity had a substantially elevated risk of developing severe disease (**Figure 7I**). Central obesity was associated with a 33.3% risk of DHF compared to 25.4% in patients without central obesity, representing an absolute risk increase of 7.9 percentage points and a relative risk of 1.22 (95% CI: 1.01-1.46). These clinical findings support our mechanistic discoveries regarding SOAT-mediated LD remodelling. The increased severe dengue susceptibility in obese patients is consistent with our demonstration that flaviviruses exploit CE-enriched lipid droplets for morphogenesis.

## Discussion

This study establishes cholesteryl ester synthesis as key host dependency for flavivirus infection and provides mechanistic insights into how viruses exploit LD metabolism to support their replication and assembly. Our findings demonstrate that SOAT1/2-mediated cholesterol esterification generates specialised LDs with liquid crystalline phases that serve as essential metabolic hubs for viral morphogenesis, advancing our understanding of flavivirus-host interactions.

The dramatic ∼100-fold reduction in viral production upon SOAT1/2 deficiency, coupled with the complete absence of assembled virions, underscores the absolute requirement for cholesterol esterification in flavivirus morphogenesis. Importantly, our findings reveal specificity for cholesterol esterification over triacylglycerol synthesis, suggesting that flaviviruses have evolved to preferentially exploit cholesteryl ester-enriched LDs rather than conventional triacylglycerol-rich LDs.

Our proteomic analyses reveal that flavivirus infection transforms LDs into specialised organelles with fundamentally altered composition and function. The enrichment of fatty acid remodelling enzymes, Rab-GTPases, sphingomyelinases, and lipid transport proteins establishes these remodelled LDs as metabolically active hubs rather than passive lipid storage depots. The recruitment of bridge-like lipid transfer proteins, and correlation with TMEM41B expression provides a potential molecular mechanism for directional lipid transfer from CE-LDs to viral replication and assembly sites. This finding links previous observations of TMEM41B’s requirement for flavivirus replication with the broader context of virus-induced lipid mobilisation^43^.

The direct interactions between cholesterol and viral proteins prM, capsid, and NS1, which specifically depend on LD-derived cholesteryl esters rather than free cholesterol, demonstrate that the cellular source of cholesterol critically determines its functional role in viral morphogenesis. The failure of exogenous free cholesterol supplementation to rescue viral production in SOAT-inhibited cells, and its apparent detrimental effects, reinforces the specificity of this dependency. The interaction of cholesterol with both structural proteins (prM, capsid) and the secreted glycoprotein NS1 suggests multiple roles for cholesterol throughout the viral lifecycle. Our work also highlights the importance of organelle contact sites, particularly ER-LD contacts, as potential platforms for virus-host interactions and therapeutic targeting. The temporal coordination between LD accumulation, turnover, and viral replication phases may represent a programmed sequence optimised to provide lipid building blocks precisely when needed for replication organelle biogenesis and virion assembly.

The clinical relevance of our mechanistic findings is supported by analysis of dengue patients from a Sri Lankan cohort, where adiposity showed significantly higher rates of severe dengue haemorrhagic fever compared to lean patients. This observation aligns with our demonstration that flaviviruses exploit cholesteryl ester-enriched LD, as obesity involves altered lipid metabolism and increased LD abundance that may provide enhanced cellular environments supporting viral replication.

Our findings contribute to the growing recognition that viruses extensively reprogramme host lipid metabolism to support their replication^15–17^. Future studies will examine whether this dependency extends to other flaviviruses and related RNA viruses, and whether these mechanisms can be captured in patient samples. The development of more potent and selective SOAT inhibitors will be essential for advancing therapeutic applications targeting host lipid metabolism to combat flavivirus infections.

## Acknowledgements

The authors acknowledge the Wellcome Trust (grants 220776/Z/20/Z and 223107/Z/21/Z to SS), BBSRC (BB/Y000307/1 to SS), an MRC studentship (AB-K) and the NIH, USA (grant number 5U01AI151788-02 to GNM).

## STAR Methods

### Cell Culture and Viral Infections

Huh7 cells were maintained in Dulbecco’s Modified Eagle Medium (DMEM) high glucose (Sigma-Aldrich, D6546) supplemented with 10% foetal bovine serum (FBS) at 37°C with 5% COL. Zika virus (ZIKV, strains MR766, NC-14-5132) and Dengue virus (DENV, strain 16681) were propagated in Vero or C6/36 mosquito cells and titrated by plaque assay on Vero cells as previously described^6,48^. The Vero E6 cell line was maintained in DMEM high glucose with 1% FBS. All cell lines were maintained at 37°C with 5% CO_2_.

For infection experiments, Huh7 cells were seeded at a density of 1×10^5^ cells per well in a 12 well plate. The following day, Huh7 cells were washed 1x with PBS and infected with ZIKV MR766 at a multiplicity of infection (MOI) of 2 in DMEM containing no FBS. 2 hours later, cells were washed 2x with PBS and replaced with DMEM containing 10% FBS. 24 and 48 hours post infection, the supernatant was collected and stored at -70°C for plaque assays. Cells were washed 1x with PBS before collecting cells for western blotting and qRT-PCR. For immunofluorescence, cells were washed 2x with PBS before fixation.

### Generation of Knockout Cell Lines

Guide RNAs (gRNAs) were designed using CCtop^49^. CRISPR RNAs (crRNAs) and trans-activating CRISPR-RNAs (tracrRNAs) were annealed by heating to 95°C followed by slowly cooling to room temperature to form crRNA/tracrRNA duplexes. Exonic and intronic targeting duplexes were combined in a 1:1 ratio and complexed with HiFi Cas9 Nuclease V3 (Integrated DNA Technologies, 1081060) at a final concentration of 44 µM crRNA/tracrRNA hybrids and 36 µM Cas9 to generate ribonucleoprotein (RNP) complexes. 1 µl of ribonuclease proteins was mixed with Huh7 (200,000 cells total) and electroporated using the Neon Transfection System (MPK10096; HiTrans 1400V 20 ms width, one pulse). Electroporated cells were cultured to establish CRISPR-edited cell pools, and genomic DNA was subsequently extracted using the DNeasy kit (Qiagen, 69504) for genotyping analysis. Pooled knockouts were single cell sorted using flow cytometry into 96 well plates containing DMEM with 20% FBS.

### siRNA Transfection

Small interfering RNAs (siRNAs) targeting MBOAT and ZDHHC family members were obtained from Dharmacon (ON-TARGETplus SMARTpool). Huh7 cells were transfected with 25 nM siRNA using Lipofectamine RNAiMAX (Thermo Fisher Scientific) according to manufacturer’s protocol. Scrambled siRNA was used as negative control. Knockdown efficiency was verified by qRT-PCR 48 hours post-transfection.

### Chemical Inhibitors and Treatments

Inhibitors were added 2 hours after Zika virus infection in complete DMEM (+10% FBS). To inhibit SOAT1/2, avasimibe (Cayman Chemical, 18129) was used at a final concentration of 20 µM. DGAT1 and DGAT2 were inhibited with 80 µM PF-04620110 (Cayman Chemical, 16425) and 40 µM PF-06424439 (Cayman Chemical, 17680), respectively.

Oleic acid (Sigma-Aldrich, O1008) or cholesterol (Sigma-Aldrich, C4951) was added 2 hours or 24 hours after Zika virus infection in combination with DMSO or SOATi in DMEM + 10% FBS. A final concentration of 500 µM oleic acid in 0.6% bovine serum albumin (BSA) was used (stock 10mM oleic acid in 12% BSA). Cholesterol (Sigma-Aldrich, C4951) was used at a final concentration of 0.1mM.

### Quantitative RT-PCR

Huh7 cells were washed 1x with PBS and then lysed with 350 µl RLT buffer (Qiagen, 74104) + 1% beta-mercaptoethanol for 30 min before storing at -70°C. RNA was isolated using the RNeasy kit (Qiagen, 74104) and subsequently used for qRT-PCR using the One-Step SYBR Green PrimeScript RT-PCR kit (Takara, RR066B).

**Table 1.**
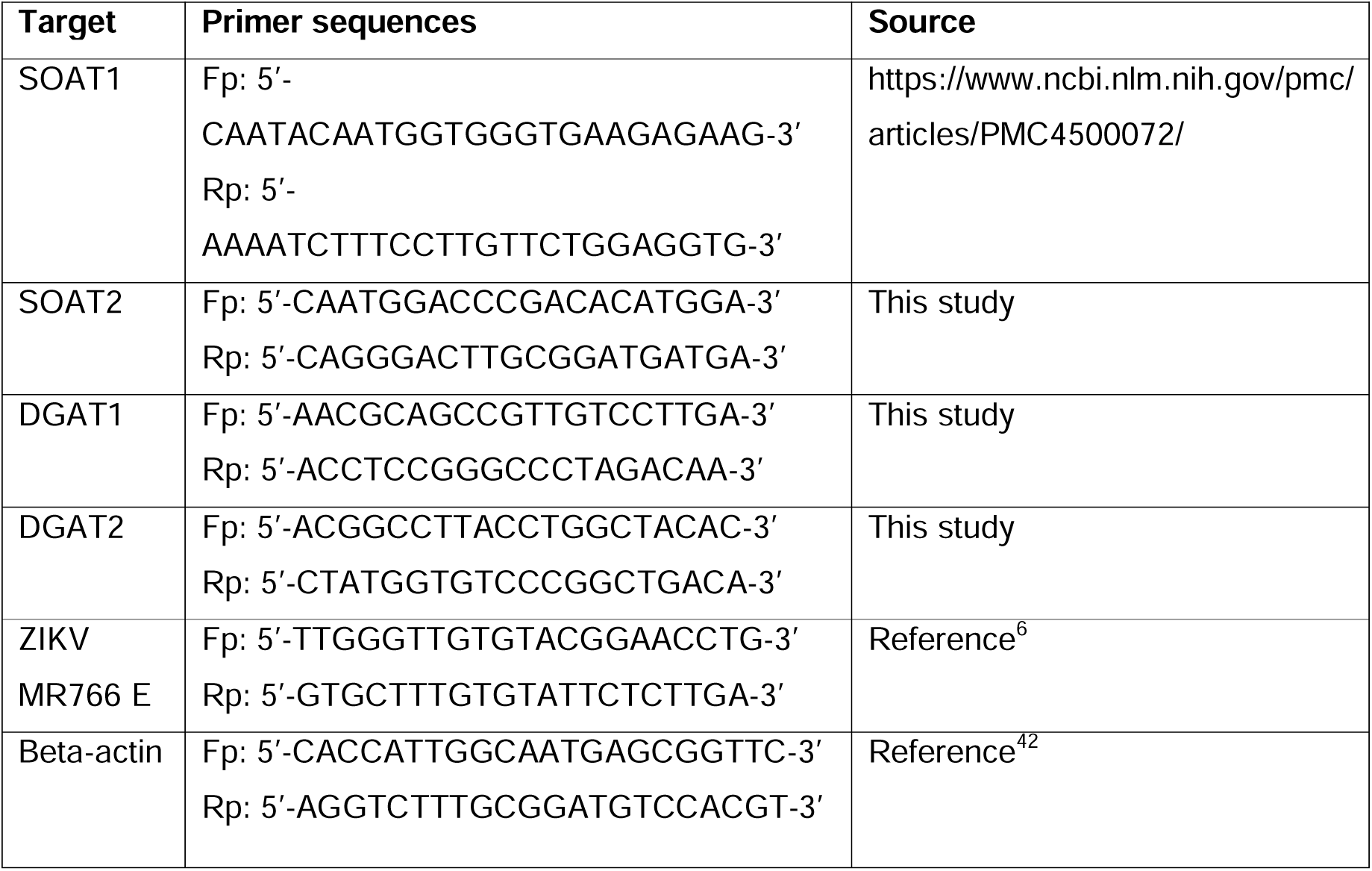
Primer sequences for qRT-PCR.

### Viral Titration Assays

Supernatant from ZIKV infections were thawed on ice and 10-fold diluted from 10^-1^ to 10^-4^ in DMEM. 100 μl of each dilution was added to a 24-well plate in triplicate, and 2.5 x 10^5^ Vero cells in 500 μl DMEM + 1% FBS were added to each well. Cells were incubated at 37°C for 2 hours, before adding 500 μl 1.5% carboxymethylcellulose sodium salt (Sigma-Aldrich, C5678) in DMEM with 5% FBS. After incubation at 37°C for 3.5 days, cells were washed 1x with PBS before adding amido black stain (10 mM Naphthol Blue Black (Sigma-Aldrich, 195243), 1 M glacial acetic acid, 170 mM sodium acetate) for 1 hour at RT.

### Immunofluorescence Microscopy

Huh7 cells were grown on 19 mm coverslips in 12 well plates for immunofluorescence staining. After Zika virus infection, Huh7 cells were washed 2x with PBS, fixed with 4% paraformaldehyde (Thermo Fisher, 43368) in PBS for 15 min at RT, washed 3x with PBS, and permeabilised with 0.1% Triton in PBS for 10 min at RT. Cells were then washed 1x with PBS before blocking with 10% normal goat serum (NGS) (Thermo Fisher, 31873) in PBS for 1 hour at RT. Cells were then incubated with primary antibodies in 5% NGS in PBS overnight at 4°C. The following day, cells were washed 4× with PBS for 5 min each and incubated with the appropriate secondary antibodies and BODIPY 493/503 (Invitrogen, D3922, 1:500) in 5% NGS in PBS for 1 hour at RT. Cells were washed a further 4× with PBS for 5 min each and then mounted onto glass slides with mounting media containing DAPI (Abcam, ab104139). The following primary antibodies were used: rabbit ZIKV E (GeneTex, GTX133314, 1:200) and mouse dsRNA (Sigma-Aldrich, MABE1134, 1:100). The secondary antibodies used were Alexa Fluor 555 anti-mouse IgG (Invitrogen, A-21422, 1:500) and Alexa Fluor 647 anti-rabbit IgG (Invitrogen, A-21244, 1:500). Stained cells were imaged using a Zeiss 880 Airyscan confocal microscope at a magnification of 63x with oil immersion (NA: 1.40).

### Live imaging of ZIKV-infected Huh7 cells

Huh7 cells were seeded at a density of 5×10^4^ cells per well in ibidi 24 well plates (cat no 82426) were treated for 1hr with with 10 μM verapamil, a broad-spectrum efflux pump inhibitor, and stained with 100nm SiR-Tubulin and SPY-555 DNA (Spirochrome, SC002, SC201). 24hrs post-seeding and three hours prior to commencing imaging, cells were washed with 1× PBS, and virus suspension equivalent to 5 and 10 MOI was applied to the cells and incubated for 1 h on ice. Post-adsorption, the cells were washed thrice with 1XPBS to remove unbound virions, and the lipid droplets were stained using the BODIPY-493/503 (Invitrogen) dye at a concentration of 1:5000. The cells continued to be incubated with SiR-tubulin and SPY-555 DNA staining solution throughout imaging.

Live-cell imaging was performed on an Evident FV4000 point scanning confocal microscope, using an Evident UPLSAPO 60x 1.3 NA silicone objective lens, with a scan area of 1024×1024 pixels at 1.98x zoom, with a pixel size of 0.105um. Images were acquired with a resonant scanner in single trip mode with 5 x frame averaging, and a pinhole diameter of 255um, on to SilVIR detectors. 13 μm Z stacks with z-spacing of 1 μm were acquired at 8 individual positions per condition, see supplementary for channel details Time-lapse images were acquired every 30 minutes over a total time course of 24 hours. Acquisition was performed in Evident CellSens Fluoview software, version 3.2.1.85.

### Image analysis and quantification

Lipid droplet identification and counting was performed using an automated pipeline created in arivis (ZEISS, https://www.zeiss.com/microscopy/en/products/software/advanced-image-analysis.html). First, the Cellpose (Stringer, C., Wang, T., Michaelos, M. et al. Cellpose: a generalist algorithm for cellular segmentation. Nat Methods 18, 100–106 (2021). https://doi.org/10.1038/s41592-020-01018-x) algorithm using the Cyto2 model was used to segment nuclei, and the Blob Finder tool was used to segment lipid droplets. Objects were size filtered to exclude small artifacts less than a volume of 15µm3. The compartments operation was used to count lipid droplets per cell. Cell tracking over time was performed using the segmented nuclei objects, using the Brownian motion type and a maximum distance of 70 μm from the nucleus centroid.

### Transmission Electron Microscopy

Cells were seeded on 13 mm coverslips in 12 well plates. After 24 hours of Zika virus infection with an MOI 2, the supernatant was removed and pre-warmed fixative agent (4% formaldehyde, 2.5% glutaraldehyde, 0.1 M PIPES buffer) was immediately added to cells for 1 hour at RT. Samples were then stored at 4°C until processing.

Ultrathin sections (90 nm) were cut using a Leica UC7 ultramicrotome, post-stained with lead citrate and imaged on a Thermo Fisher Tecnai T12 TEM at 120 keV (with Gatan OneView CMOS camera).

#### Cryo-FIB milling and cryo-TEM data acquisition

Lamellae were prepared from Huh7 cells grown on gold Quantifoil R2/1 and plunge frozen in liquid ethane, using the Arctis (ThermoFisher Scientific) plasma FIB. Prior to milling, the cells were coated with platinum followed by a layer of organoplatinum (trimethyl(methylcyclopentadienyl)platinum(IV)) using the gas injection system. Selected cells were successively thinned using a 30kV plasma beam and currents 4nA to 0.1nA before being polished using a current of 30pA.

Tilt series data was collected on Titan Krios (ThermoFisher) cryo-electron microscope equipped with a Falcon 4i detector and a SelectrisX energy filter. Data acquisition was performed using Pacetomo^50^ scripts in SerialEM^51^ covering an angular range of ±60° in 3° increments using a dose-symmetric scheme^52^. Each tilt series was collected with a nominal defocus of 5µm and movies were recorded in EER format^53^ using a dose of 2.91 e/Å^2^ per tilt resulting in a cumulative dose of 120 e/Å^2^ over the full tilt series. Magnification was nominally 64000x resulting in a calibrated pixel size of 1.97 Å/px on the detector.

#### Tomogram reconstruction

WarpTools^54^ was used to perform reference-free motion correction, CTF and defocus estimation on the individual frameseries data followed by the generation of dose-weighted tilt-series files. Tilt series were then aligned using AreTomo2^55^ and the alignments were imported back into WarpTools for generating binned CTF-corrected tomograms as well as half-sets for denoising to boost contrast and aid the interpretability of the tomograms.

### Lipid Droplet Isolation

Lipid droplets were isolated by sucrose density gradient centrifugation as previously described. Briefly, cells were homogenised in buffer A (20 mM Tricine-KOH pH 7.8, 250 mM sucrose, 1 mM EDTA) using a Potter-Elvehjem homogenizer. The homogenate was adjusted to 20% sucrose and overlaid with 5% sucrose in buffer A. After centrifugation at 152,000 × g for 1 hour at 4°C, the LD fraction was collected from the top interface.

### Proteomics Analysis

Isolated lipid droplets were lysed in 8 M urea buffer, proteins were reduced with dithiothreitol, alkylated with iodoacetamide, and digested with trypsin. Peptides were analysed by liquid chromatography-tandem mass spectrometry (LC-MS/MS) using an Orbitrap Fusion Lumos mass spectrometer. Data were processed using MaxQuant software and analysed using Perseus for statistical analysis and volcano plot generation.

### Virus-Like Particle (VLP) Assembly Assay

1 µg of a pcDNA plasmid containing ZIKV prM and E [11] was transfected into Huh7 WT, SOAT1 and SOAT2 knockout cells using Lipofectamine 3000 (Invitrogen, L3000015). Cell lysates were collected 48 hours in NP-40 cell lysis buffer (50 mM Tris-HCl pH 7.5, 150 mM NaCl, 2 mM MgCl2, 5 mM EDTA, 0.5% NP-40) with protease inhibitors after transfection.

### Acylation Profiling

For lipid acylation analysis, cells were labelled with [¹LC]-oleic acid (50 μCi/ml) for 4 hours. Lipids were extracted using chloroform:methanol (2:1) and separated by thin-layer chromatography (TLC) on silica gel plates. Radioactive signals were detected by autoradiography.

For protein acylation analysis, cells were labelled with alkyne-C16 (50 μM) for 4 hours, lysed, and subjected to click chemistry using azide-biotin. Biotinylated proteins were captured using streptavidin beads and analysed by Western blot.

### Cholesterol tracking

For cholesterol tracking experiments, cells were incubated with alkyne-cholesterol analogue (10 μM) for 16 hours before infection. At indicated time points, cells were fixed and subjected to copper-catalyzed azide-alkyne cycloaddition (CuAAC) click chemistry using azide-Alexa Fluor 594 to visualize cholesterol distribution.

For photo-crosslinking experiments, cells were labelled with diazirine-alkyne cholesterol analogue (10 μM) for 16 hours, infected with ZIKV, and exposed to UV light (365 nm, 15 minutes) at 24 hours post-infection. Crosslinked proteins were captured using CuAAC click chemistry with azide-biotin followed by streptavidin pulldown and Western blot analysis.

### Clinical Data Analysis

Dengue patient data from a Sri Lankan cohort (n=561) were analysed with appropriate ethical approval. Central obesity was defined as waist circumference ≥80 cm in women or ≥90 cm in men. Disease severity was classified as dengue fever (DF) or dengue haemorrhagic fever (DHF) according to WHO criteria. Statistical analysis was performed using Fisher’s exact test for categorical variables.

### Statistical Analysis

Data are presented as mean ± standard error of the mean (SEM) from at least three independent experiments unless otherwise stated. Statistical significance was determined using unpaired Student’s t-test for two-group comparisons or one-way ANOVA with Dunnet’s post-hoc test for multiple comparisons. P-values <0.05 were considered statistically significant. Statistical analyses were performed using GraphPad Prism 9 software.

