## Supplemental information for "Remodelled cholesteryl ester enriched lipid droplets fuel flavivirus morphogenesis"

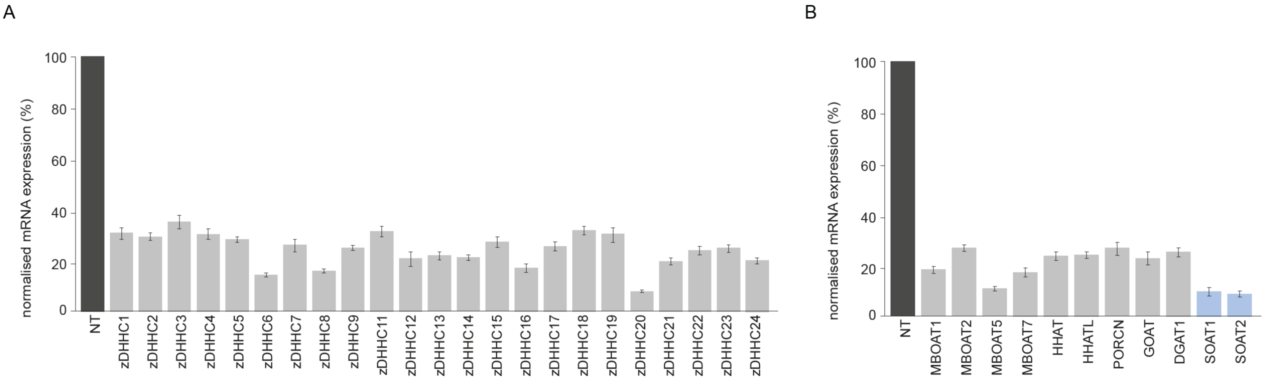


**Figure S1: siRNA screen of zDHHC and MBOAT acyltransferases**

**(A)** Normalised mRNA expression levels of zDHHC family genes following siRNA-mediated knockdown in Huh7 cells. NT, non-targeting control siRNA. **(B)** Normalised mRNA expression levels of MBOAT family genes following siRNA-mediated knockdown in Huh7 cells. Data are presented as mean ± SEM from three independent experiments. mRNA levels were quantified by RT-qPCR and normalised to the non-targeting control (set to 100%).

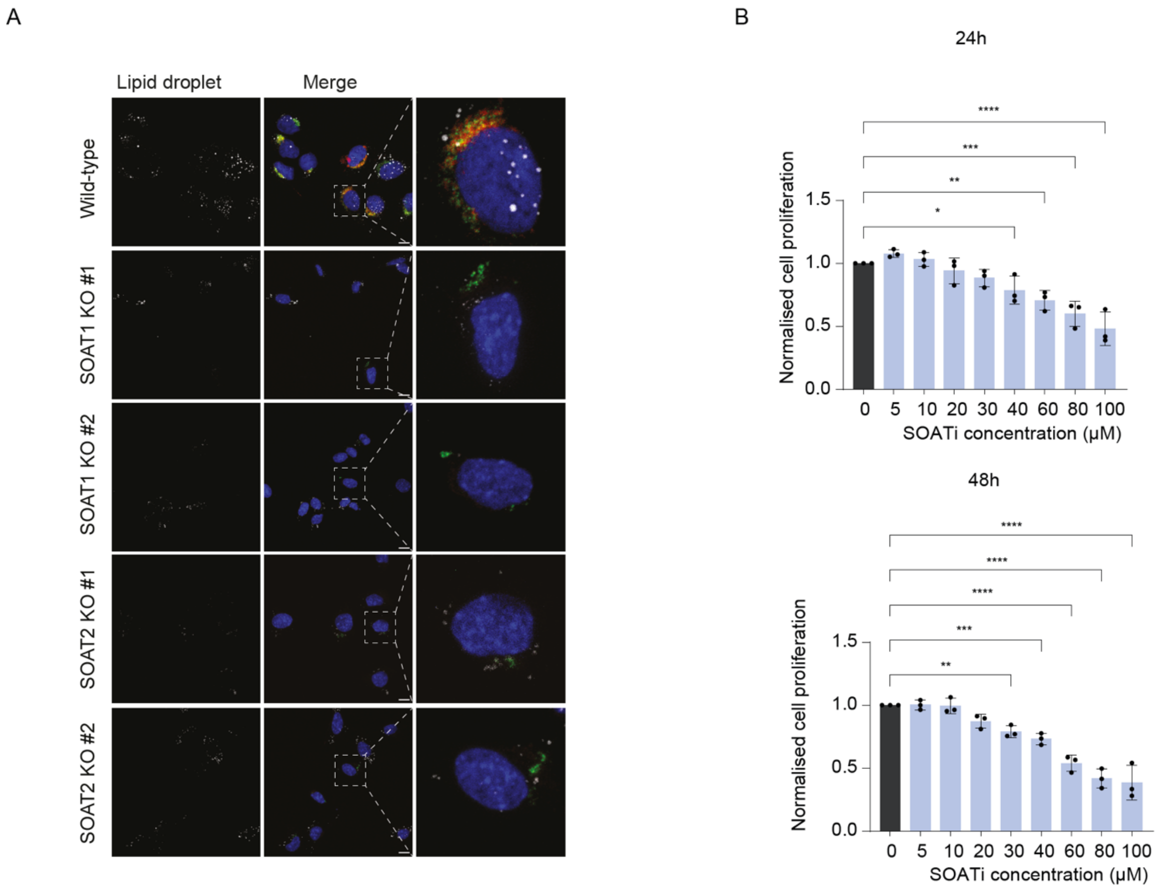


**Figure S2: SOAT deficiency reduces lipid droplet abundance**

**(A)** Confocal microscopy images of lipid droplets in wild-type and SOAT1/2 knockout cell lines. Lipid droplets were stained with BODIPY 493/503 (white), nuclei with DAPI (blue). Scale bar = 10 μm. Dashed boxes indicate regions shown at higher magnification in the right panels. Lipid droplet numbers are reduced in SOAT1 and SOAT2 knockout cells compared to wild-type cells. **(B)** Cell proliferation assay showing dose-dependent effects of SOAT inhibitor (SOATi) treatment at 24h and 48h. Huh7 cells were treated with increasing concentrations of Avasimibe (0-100 μM) and cell proliferation was measured. Data are presented as mean ± SEM from three independent experiments, normalised to untreated control. Statistical significance was determined by one-way ANOVA with Dunnett's multiple comparisons test (*p<0.05, **p<0.01, ***p<0.001, ****p<0.0001).

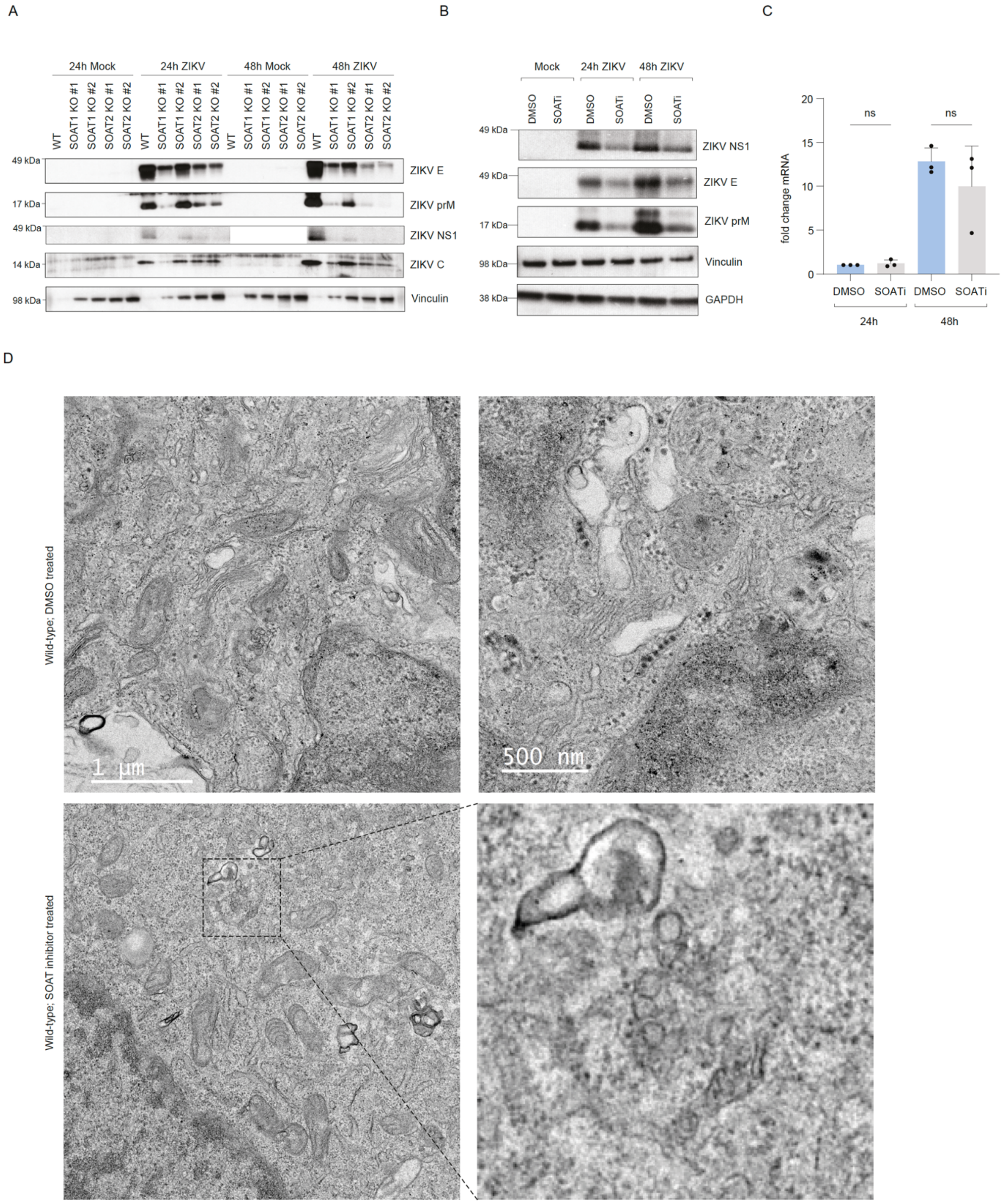


**Figure S3: SOAT deficiency reduces viral protein levels and impair replication organelle formation.**

**(A)** Western blot analysis of viral proteins in wild-type, SOAT1$^{-/-}$, and SOAT2$^{-/-}$ cells at 24 and 48 hours post-infection with ZIKV. Vinculin serves as loading control. Viral proteins (E, prM, NS1, capsid) are reduced in SOAT-deficient cells compared to wild-type. **(B)** Western blot analysis of viral proteins in wild-type cells treated with DMSO (control) or SOAT inhibitor (SOATi) at 24 and 48 hours post-ZIKV infection. GAPDH and Vinculin serve as loading controls. **(C)** Quantitative RT-

PCR analysis of viral RNA levels in DMSO- versus SOATi-treated cells at 24 and 48 hours post-infection. Data are presented as fold change relative to DMSO control (mean ± SEM, n=3 independent experiments). ns, not significant. **(D)** Transmission electron microscopy images of wild-type cells treated with DMSO (top panels) or SOAT inhibitor (bottom panels) at 24 hours post-ZIKV infection. Right panels show higher magnification views.
